# Predicting epitopes Based on TCR sequence using an embedding deep neural network artificial intelligence approach

**DOI:** 10.1101/2021.08.11.455918

**Authors:** Norwin Kubick, Pavel Klimovich, Mariusz Sacharczuk, Michel-Edwar Mickael

## Abstract

T cells receptors are fundamental in recognizing antigens and mediating an appropriate specific immune response against them. Today TCR sequencing has contributed to forming a large repertoire for different immune-associated pathologies. However, predicting epitopes based on TCR sequences has not been satisfactory achieved. We formed a deep neural network using a combination of an embedding autoencoder and selu and relu layers to predict epitopes based on TCR TCR β-chain CDR3. We trained our model using the VDJ database (VDJdb) and validated it using the manually curated catalog of pathology-associated T cell receptor sequences (McPAS-TCR). We used various metrics to measure the accuracy of our tool. We found that our tool can achieve an accuracy of 98 %. Overall our approach presents a step toward identifying microbes crosslinking epitopes that could be playing an important role in various immune diseases.

## Introduction

TCR sequencing is increasingly becoming a crucial technique in adaptive immunity research. One of TCR sequencing applications is investigating the pathogenesis of diseases associated with T cells. For example it has been shown that CDR3 clono-type TCR cross reaction to drugs could cause severe adverse reactions [1]. Additionally TCR sequencing is used in monitoring the immune response to infectious diseases or vaccines, where it could be used to analyses the dynamics of appearance of new clono-types that are specific for the virus or the vaccine being studied and their status transfer from naïve to memory [2]. Furthermore, TCR sequencing could also be used in Facilitating diagnosis and early recurrence detection. It was demonstrated that CDR3 could be used to distinguish between different stages of cutaneous T-cell lymphoma.

TCR variation is due to various factors. Individual T cells expresses a distinctive T-cell receptor (TCR). Gene recombination of the VDJ gene regions results in constructing unique α and β chain that form the TCR. Unique TCRs are capable of recognizing a unique group of epitopes presented to it by the MHC expressed on antigen presenting cells. The uniqueness of each TCR is based on the CDR1, CDR2, CDR3 regions. Of them the most crucial regions was shown to be CDR3. There are also other factor that contribute to TCR specificity such as the type of CD4+ T cells. CD4+ T cells are a heterogonous group of cells with pro and anti-inflammatory functions [3], [4] [5]. For example that Th1 and Th17 could be pro-inflammatory, while Th2 and Tregs could be anti-inflammatory. TFH play an important role in initiating an immune response through activating B cells. CD4+ T cells also differ due to their interlukins environment, where they can change from one phenotype to other[6]. Furthermore, the difference could be also as specific as a single cells and difference in epigenetic activation [7].

One way to address the heterogeneity in CD4+ T cells and hence thee variation in their TCR repertoire is differentiating between these cells based on their epitope sequence. However, Different experimental strategies are currently available to determine epitope-specific TCRs, i.e., TCRs that recognize the epitope of interest, such as epitope-MHC multimer assays and peptide stimulation experiments. However, these techniques are not error-free as they can miss important epitope-specific TCRs or falsely report non-binding TCRs to be epitope specific (8). Nevertheless, these methods have been extensively used to generate a large amount of epitope-specific TCR data which has been collected in public databases such as the VDJdb and McPAS-TCR [8].Computational techniques based on either machine learning and AI also exist. However they suffer from drawbacks. Machine learning techniques are usually less accurate than AI approaches. One of the important AI techniques is RGO (pEptide tcR matchinG predictiOn)[9]. However ERGO suffer from drawback of using hot-one encoder. The main disadvantage, of using hot-one encoder, is the unawareness of the model of the inherent relationship between the TCR sequence and the epitope.

In this report, we used a AI neural network to predict epitopes based on TCR sequence. We imported V database and used it as the training model for our AI workflow. In the first step of the AI workflow, we used a google universal encoder to embed the sequence each CDR3 TCR sequence and its respective epitope into a matrix of (1,512). Then we a deep network consisting of two main layers in kereas; (i) selu and (ii) relu. After that the predicted sequence was decoded using a sigmoid function and a cosine distance was used to predict the nearest epitope. Overall the accuracy of the model is more than 98% and the loss is less than 0.005.

## Methods

### Compilation of TCR Dataset

For construction of our network we employed the VDJdb database. This database is a carefully selected database of T-cell receptor (TCR) sequences with identified antigen specificities. The database includes the following entries; Gene which represents TRA or TRB (comprising either TCR chains alpha and beta chins respectively). CDR3; represents the CDR3 amino acid sequence, V(variable sequence segment allele), J(joining segment allele), species(humans, chimps), MHC chain alleles(A,B), MHC class (I,II), epitope, epitope gene, epitope species. We only selected CDR3 sequence and their relevant epitopes, regardless MHC class or species.

#### Description of the workflow

Then we employed a universal google encoder to embed the sequences[10][11][12] After that we divided the resulting matrices in KERAS for two groups (i) test (ii) validation. Next we applied a deep neural network to classify the epitope matrices based on the TCR sequences. The model consisted of sequential KERAS layers SELU and RELU. RELU function was shown to perform better than other r activation function as it does not activate all the neurons at the same time. The SELU (Scaled Exponential Linear Unit), is used to combine the effects of the RELU and the batch normalization. Finally we used our model to predict novel sequences based on the cosine distance between the predicted matrix and the epitope embedding.

Evaluation of the model was done using two methods (i) first we used the validation function in KERAS, where the data was divided into two groups(training) and test. (ii) we decoded the prediction of our model and computed the nearest value using cosine distance. To achieve the highest accuracy, We used several loss functions and selected the one with the highest accuracy

## Results

### The workflow is composed of five major steps

First we downloaded the CDR3 sequence and th epitopes. Then we employed a universal google encoder to embed the sequences. After that we divided the resulting matrices in KERAS for two groups (i) test (ii) validation. Next we applied a deep neural network to classify the epitope matrices based on the TCR sequences. Finally we used our model to predict novel sequences based on the cosine distance between the predicted matrix and the epitop embedding.

### Model performance

Our model can predict the TCR –epitope relationship with an accuracy of 98% (figure 2a). This accuracy is achieved after 5 epochs (figure 2a). The model loss achieved was less than 0.1 % using the MeanSquaredLogarithmicError loss function in KERAS. Our results indicate that the highest accurac and the lowest loss could be achieved using this particular loss function. Employing other loss functions resulted in either lower accuracy or higher loss (Table 1).

**Figure 1.**
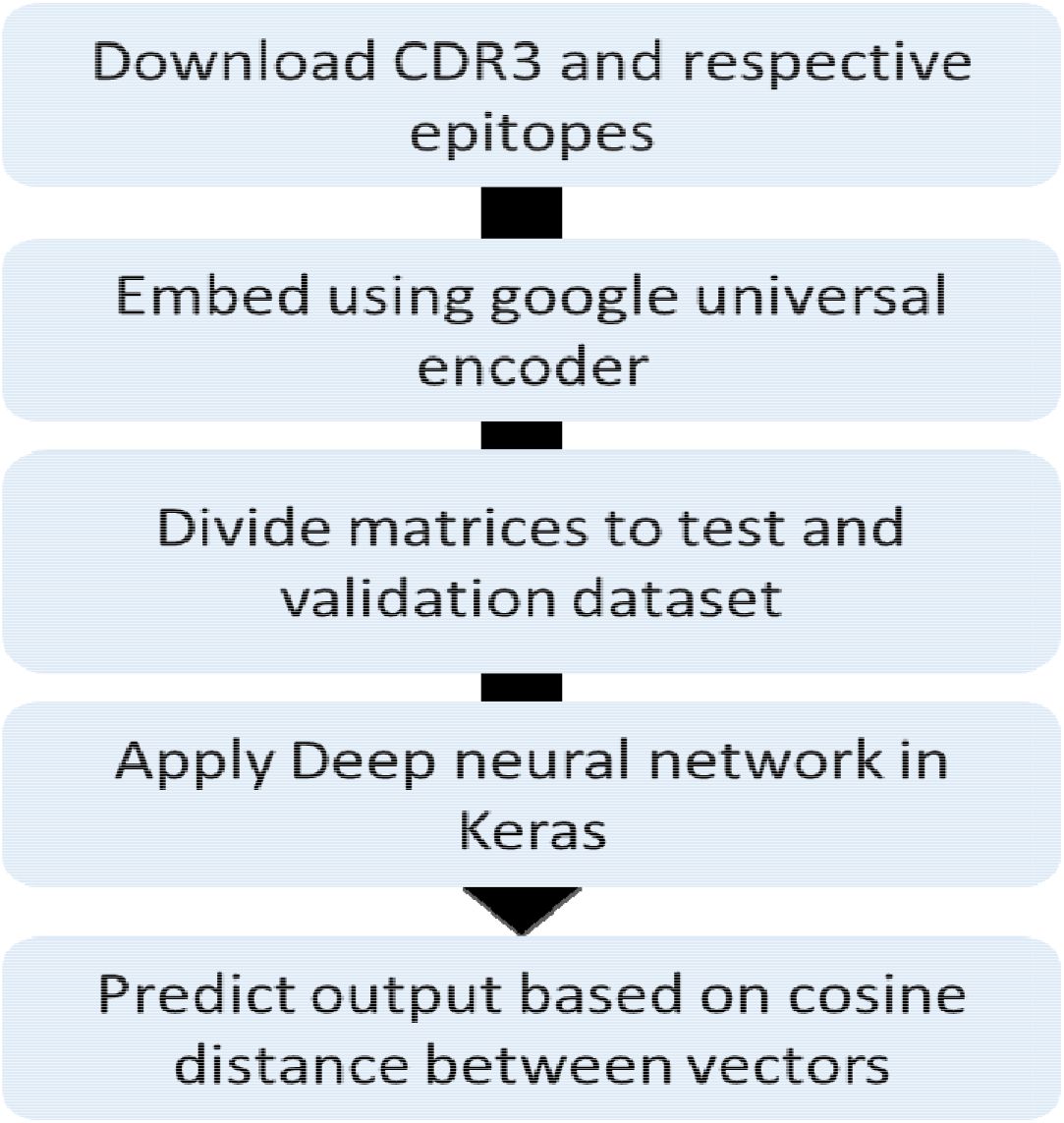
Model workflow consists of several main step including preparing the data, embedding, preprocessing, applying the neural network and prediction.

**Figure 2.**
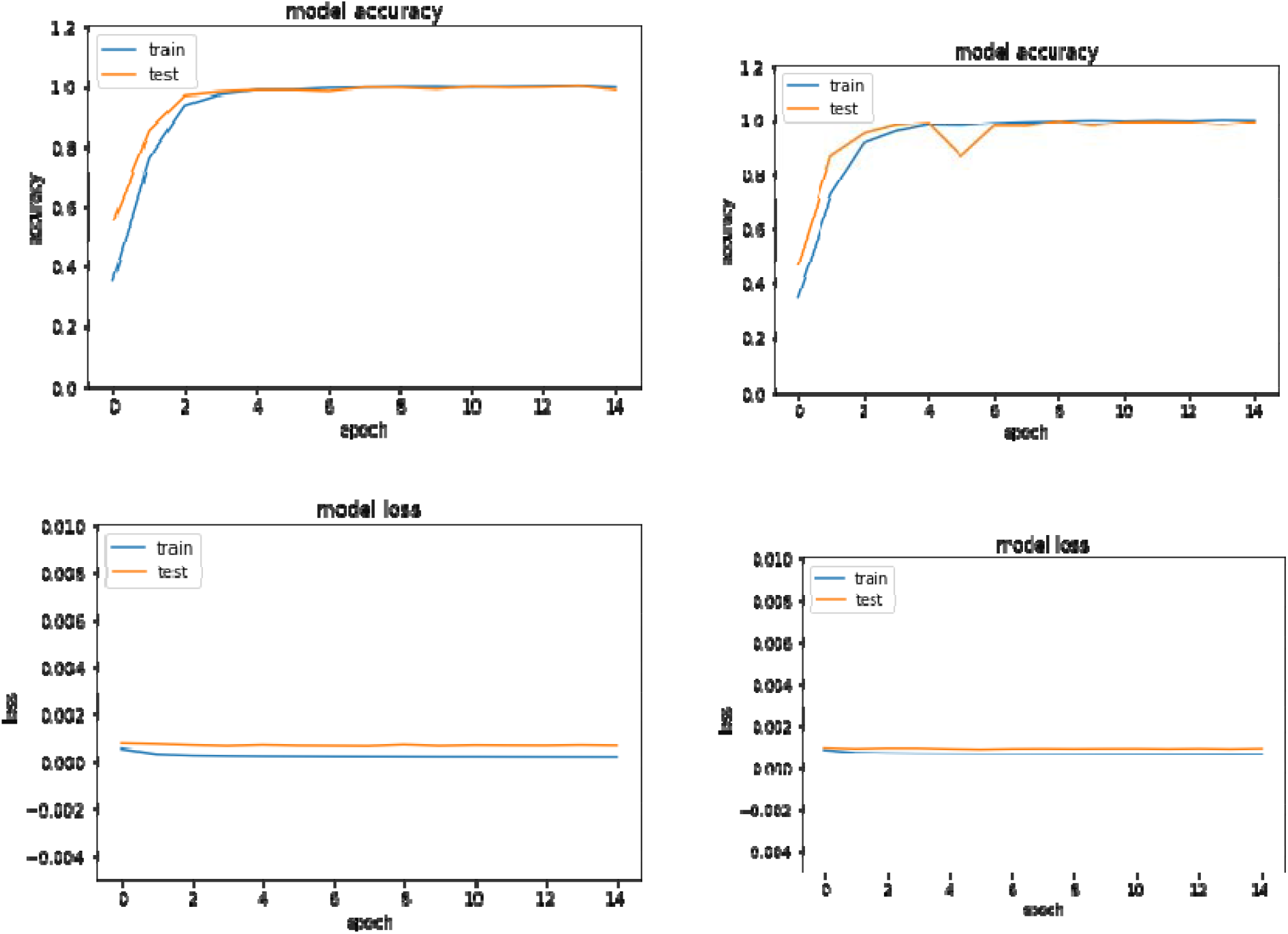
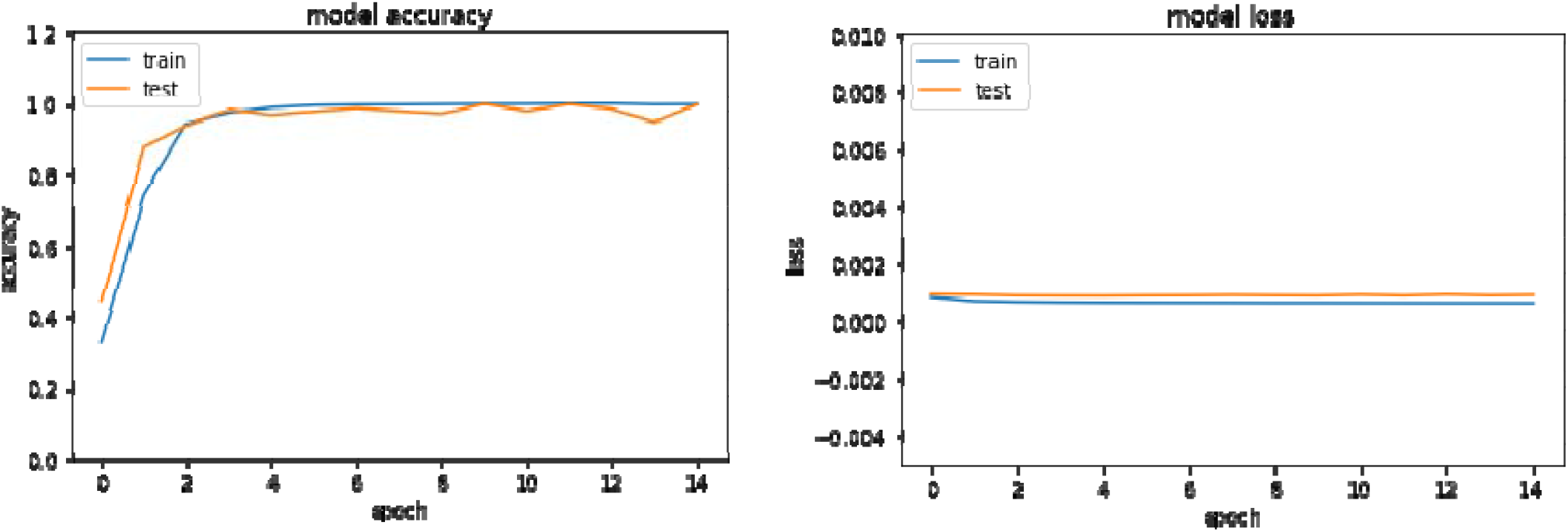
Our model achieves 98 % accuracy and zero loss. **A1)** MeanSquaredLogarithmicError accuracy a2) MeanSquaredLogarithmicError loss b1) Huber accuracy b2) huber loss c1) LogCosh accuracy c2) LogCosh loss.

| Function | Accuracy | Loss |
| --- | --- | --- |
| MeanSquaredLogarithmicError | 98-99% | <0.01 |
| categorical_crossentropy | 99% | >50 |
| BinaryCrossentropy | <1 | <0.1 |
| CategoricalHinge: | <1 | <0.1 |
| CosineSimilarity | 98-99% | <0.3 |
| Hinge: | <1 | <0.9 |
| huber | 98-99% | <0.01 |
| KLDivergence: | <1 | <0.1 |
| LogCosh | 98-99% | <0.01 |

## Discussion

### Model performance

Our model outperform existing models. Epitope prediction is normally based on using TCR sequence, and Deep learning to predict the epitope sequence. This problem has been tackled by various groups [9][13][14] [8]. The training dataset for this network consists of CDR3 TCR in addition to known respective epitopes. The first step to tackle this problem is to convert the amino acid sequence into numerical matrices. Previously, a one-hot vector was used to represent each amino acid of the CDR3 sequence. The hot vector consisted of 20 positions with zeros, except for the position representing the amino acid. All the amino acids of any CDR3 sequence were joined and terminated with a “stop codon”. Padding using zeros was used to ensure all the sequences have equal lengths. The result was used as the input of an autoencoder network that consists of two parts as described before. The first part was an encoder, and it consisted of two layers with an exponential linear activation unit. The decoder was built in the same form as the encoder but in reverse order. The authors applied a softmax function which is responsible for translating the last layer into a decoder layer. However one-hot encoding is blind regarding the relationship between CDR3 sequence and the epitope. Another group embedded the epitope using an an LSTM (long short-term memory) network. After that, the two embedding were fed into an MLP (multi-layer perceptron) function. [9]. However MLPS suffer from three main disadvantages (i) MLP with hidden layers possess a non-convex loss function with more than one local minimum. (ii) MLP usage necessitates tuning a number of hyper parameters (iii)MLP is highly sensitive to feature scaling. Our model overcome this problem by using a deep neural networks of two main layers an SELU and RELU.

Our model is not without limitations. Although our model is capable of predicting de novo TCR sequences. The decoding procedure is based on the multiplication of a sigmoid function with the output of the deep network, Then this is followed by calculating the cosine distance to predict the relevant epitope. As discussed in the methods section, our model does not use one-hot encoding as it is aware of the structure and the specification of the epitopes. However a future step would to embed each amino acid as different vector and build epitopes as a matrices and run the new formed matrices through the our model. Another addition is the use of LSTM model to increase awareness between each epitope sequence letters. Future applications of our model would be able to predict interactions not only between TCR and epitopes but other cell membrane components such as GPCRs as well as histone modifications[15][16].

## Conclusion

In this report we introduced our deep neural network which is capable of predicting epitopes based on CDR3 with accuracy of more than 98%. The model suffers a small loss and outperform other existing model. It is also aware of the interactions between the TCR and its epitope based and avoids using one-hot encodes. However, in its current form it does not take into consideration the relationship between amino acids within the epitope.

